# Resolving intergenotypic *Striga* resistance in sorghum

**DOI:** 10.1101/2022.12.08.519579

**Authors:** Sylvia Mutinda, Fredrick M. Mobegi, Brett Hale, Olivier Dayou, Elijah Ateka, Asela Wijeratne, Susann Wicke, Emily S. Bellis, Steven Runo

## Abstract

Genetic underpinnings of host-pathogen interactions in the parasitic plant *Striga hermonthica,*a root parasitic plant that ravages cereals in sub-Saharan Africa, are unclear. We performed a comparative transcriptome study on five genotypes of sorghum exhibiting diverse resistance responses to *S. hermonthica* using weighted gene co-expression network analysis (WGCNA). We found that *S. hermonthica* elicits both basal and effector-triggered immunity – like a bona fide pathogen. Resistance response was genotype-specific. Some resistance responses followed the salicylic acid-dependent signaling pathway for systemic acquired resistance characterized by cell wall reinforcements, lignification and callose deposition while in others the WRKY-dependent signaling pathway was activated leading to a hypersensitive response (HR). In some genotypes, both modes of resistance were activated while in others, either mode dominated the resistance response. Cell-wall-based resistance was common to all sorghum genotypes but strongest in IS2814, while HR-based response was specific to N13, IS9830 and IS41724. WGCNA further allowed for pinpointing of *S. hermonthica* resistance causative genes in sorghum. Some highlights include a Glucan synthase-like 10, a pathogenesis-related thaumatin-like family, and a phosphoinositide phosphatase gene. Such candidate genes will form a good basis for subsequent functional validation and possibly future resistance breeding.

**Highlight:** Parasitic plants of the *Striga* genus are major pests to cereals in Africa. We pinpointed genetic causes of *Striga* resistance in sorghum that can be harnessed for future resistance breeding.

## Introduction

Sorghum [*Sorghum bicolor* (L) Moench] is a staple cereal for millions in sub-Saharan Africa (SSA) but, its production is greatly constrained by the parasitic plant *Striga* spp. (Orobanchaceae family). *Striga* attaches to roots of host crops to siphon water and nutrients. In sorghum, *Striga* leads to crop losses of between 30 and 100 % (Ejeta, 2007) millions of families in Africa that depend on the cereal to hunger and loss of livelihoods.

*Striga* is difficult to control because of its well-adapted lifestyle (Runo and Kuria, 2018). Parasitism is tightly cued to the host allowing seed germination only in response to host-derived biomolecules, hormones – mainly strigolactones (Matusova *et al*., 2005; Bouwmeester *et al*., 2007; Gomez-Roldan *et al*., 2008; López-Ráez *et al*., 2011). Germination is followed by the parasite attaching to the host and developing vascular vessels that create a continuum for water and nutrient acquisition (Dörr, 1997). Parasitism continues for 30-40 days below-ground after which *Striga* emerges from soil and becomes photosynthetic. *Striga* then flowers to complete its lifecycle by producing approximately 50, 000 seeds per flower (Berner *et al*.,1995).

Smallholder farmers in SSA, try to maintain low *Striga* infestation levels using agronomic and cultural practices such as hand weeding and crop rotation with non-hosts (Kanampiu *et al*., 2018) but these have achieved little or no success. A more practical and sustainable strategy would be to integrate multiple management approaches and greatly leverage natural host resistance (Mwangangi *et al*., 2021).

However, mechanics of host resistance against *Striga* are only beginning to get unraveled. At the fore are new findings that show *Striga* spp. to be sophisticated manipulators of plant cells (Li *et al*., 2009), and the parasite’s ability to trigger the host innate immune response (Su *et al*., 2020) reminiscent of host pathogen interactions (Dangl and Jones, 2006). If this model fits *Striga-host* interaction, *Striga* should firstly elicit pathogen triggered immunity (PTI) through its yet to be determined pathogen associated molecular patterns that makes it encounter mechanical barriers from the host manifesting in form of cell wall and biochemical fortifications. *Striga* should then suppress PTI and promote pathogenesis by a second offense on host through injection of effector-like molecules into plant cells (Dangl and Jones, 2006). If the host is resistant, effector triggered immunity (ETI) – whose hallmark is programmed cell death – will be activated effectively stopping further parasitism.

Responses suggestive of PTI and ETI have been reported in various sorghum genotypes. For instance, mechanical barrier-type of resistance that fortify the host cell wall to block ingression of the parasite in the east African Durra sorghum N13 (Maiti *et al*., 1984; Mbuvi *et al*., 2017; Kavuluko *et al*., 2021), programmed cell death at the interphase of the host and the parasite that is characterized by a hypersensitive response at the host-parasite junction in IS14963 (Kavuluko *et al*., 2021) and secondary metabolites such as polyphenols deposition at the interphase of the host and the parasite, in some wild sorghum accessions (Mbuvi *et al*., 2017).

What is not clear is if specific sorghum genotypes exhibit one or more modes of resistance working in concert to ward off *Striga*. So far, data favor a model in which a genotype has one – or a dominant mechanism of resistance. Genetically, such a resistance model is suggestive of multiple independent pathways activating *Striga* resistance – harbored in different genotypes. Here, we use a comparative transcriptome analysis of sorghum harboring distinct phenotypes to clarify if multiple forms of resistance can manifest in the same genotype against the alternate, that genotypes have predominantly one mode of resistance. Our findings have important implications in breeding for broad-spectrum and durable *Striga* resistance in sorghum.

## Materials and Methods

### Plant materials

We selected sorghum genotypes representing mechanical barrier resistance, hypersensitive response then used a comparative transcriptomics approach to home in on the genetic underpinnings of each resistance mechanism. Genotypes comprised, the east African Durra sorghum N13, Middle East Durra-Caudatum, IS2814 and the advanced east African Caudatum IS9830 to represent mechanical barrier resistance (Maiti *et al*., 1984; Kavuluko *et al*., 2021; Mbuvi *et al*., 2017), and IS14963, a Caudatum from Central Africa and IS41724, an Indian Durra to represent hypersensitive interactions. These sorghum materials were originally obtained from the International Crops Research Institute for Arid and Semi-arid Tropics (ICRISAT) and are maintained at the Plant Transformation Laboratory (PTL) at Kenyatta University. Seeds of *S. hermonthica* were obtained from infested farms in western Kenya Busia, Alupe (0.45°, 34.13°) in 2018. Only *S. hermonthica* seeds were used in this study.

### Conditioning and germination of *Striga* seeds

Conditioning of *Striga* seeds was done according to Mbuvi *et al.* (2017). Prior to conditioning, *Striga* seeds (25 mg) were washed in 25 ml of 10% (v/v) Sodium hypochlorite (commercial bleach) for 10 minutes with gentle agitation. Seeds were collected in a funnel lined with Whatman GFA filter paper (Meadow, UK) and thoroughly rinsed with double distilled water to get rid of Sodium hypochlorite. Seeds were then transferred on petri plates (90 mm) lined with moisturized filter papers and incubated in the dark at 30 °C for 14 days. After conditioning, the seeds were pre-germinated by adding 3 ml of 0.1 ppm GR24 (Chiralix, Nijmegen, Netherlands) and incubating overnight at 30 ^°^C.

### Sorghum germination and infection with *Striga*

Sorghum seeds were sowed in germination pots (10 x 10 x 7 cm) filled with moisturized vermiculite and watered with Long Ashton nutrient media (Hudson, 1967). Ten (10) days following germination, seedlings were transferred to rhizotrons-soil free transparent root analysis chambers of the dimensions 25 x 25 x 5 cm (Nunc, ThermoFisher Scientific, UK). The chambers were prepared as described by Mbuvi *et al.* (2017) by filling them with vermiculite and then overlaying them with 50 micron-thick nylon mesh. The chambers were then wrapped with aluminum foil and kept in a glasshouse at temperature cycles of 28 °C Day: 24 °C night with 12-h photoperiod and 60% humidity. During this period, the plants were drip-fed with nutrient media. To infect sorghum roots with *Striga*, rhizotrons were opened and sorghum roots carefully aligned with pre-germinated *Striga* seeds using a soft paint brush. After infection, the chambers were closed, wrapped in aluminum foil, and maintained in the glasshouse as described above. Three plants per genotype were screened in a randomized complete block design (RCBD). Plants were kept in a glasshouse at temperature cycles of 28 °C day and 24 °C night with 12-h photoperiods.

### Characterizing *Striga* resistance mechanisms in sorghum

Mechanisms of resistance against *Striga* were evaluated in two genotypes viz IS2814 and IS41724 using histological assays. These genotypes had previously been reported to harbor post attachment resistance based on low *Striga* number, short length, and low biomass metrics (Kavuluko *et al*., 2021). Histological analyses were done by studying the host-parasite interface, 9 days post infection (dpi) as described by Kavuluko *et al.* (2021). Briefly, tissue segments at the sorghum-*Striga* interface were dissected, fixed in Carnoy’s fixative, and stained with 1% safranin O dye. Tissues were then cleared of safranin O dye in choral hydrate solution (2.5 g/ml) overnight with gentle shaking. Following de-staining, photographs revealing the extent of parasite ingression on the host roots were documented using a Leica stereomicroscope MZ10F fitted with DFC 310FX camera.

Prior to sectioning fixed tissues were preprocessed using Technovit 7100^™^ kit following the manufacturer’s instructions (Heraeus Kulzer GmbH, Hanau, Germany) and protocols described in Mbuvi *et al.* (2017). The tissues were pre-infiltrated, (1:1, Technovit^™^ solution: 100% ethanol), infiltrated (100% Technovit^™^ solution) then embedded in 1.5 ml microcentrifuge tube lids containing Technovit^®^1 /Hardener 2 (1:15) and left to set for two weeks. Embedded tissues were mounted onto wooden blocks using the Technovit^™^ 3040 kit following the manufacturer’s instructions (HaraeusKulzer GmbH). The wooden blocks holding the tissues were set on the rotary Leica RM2145 microtome (Leica, Germany) and 5-μm thick sections were obtained. Sections were placed on hydrated glass slides, dried on a hot plate at 65 °C, stained using 0.1% toluidine blue O dye in 100 mM phosphate buffer for 2min then washed in distilled water. A drop of DePex (BDH, Poole, UK) was applied on each of the dry sections and overlaid with coverslips. The slides were observed and photographed using a Leica DM100 microscope fitted with a Leica MC190 HD camera, (both Leica, Germany).

### Lignin staining

Surface-sterilized sorghum seeds of the genotypes N13, IS41724 and IS14963 were germinated and transferred to rhizotrons. Sorghum roots were allowed to spread followed by infection with pre-germinated *Striga* seeds. At 9 dpi, root segments were excised at the point of *Striga* attachment, embedded in 7% (w/v) agar, and sectioned by a vibratome (Leica VTI200s, Leica Germany). Sections with 20-μm thickness were stained of 1% (w/v) phloroglucinol solution for 5 mins. Phloroglucinol (0.3 g) was dissolved in 10 ml absolute ethanol. This 3% phloroglucinol solution was mixed with 2M HCL in a ratio of 2:1 to make 1% for staining the sections. Photographs were taken using the Leica DM100 microscope fitted with a Leica MC190 HD camera, (Leica, Germany).

### RNA isolation, library preparation and sequencing

Root tissues were collected from sorghum infected with *Striga* at 3 and 9 days including noninfected controls. Three replicates were collected for each treatment in the five genotypes. Samples were immediately frozen and ground in liquid nitrogen. RNA isolation was carried out using the ISOLATE II RNA Plant kit (Bioline, Meridian, UK) according to the manufacturer’s recommendations. Samples were extracted in Trizol including on column desalting, DNase treatment, and purification. The quantity and quality of the isolated RNA were assessed by nanodrop and Agilent Bioanalyzer (RNA integrity Number) respectively. Samples with a RIN value of ≥7.0 were used to prepare Tag Seq libraries according to (Lohman *et al*., 2016).

### Quality control Preprocessing of raw 3’ Tag-RNA-Seq reads

The 3-Tag-RNA-Seq-analysis Next Flow pipeline (https://github.com/fmobegi/3-Tag-RNA-Seq-analysis) which wraps up the TagSeq utilities v2.0 (https://github.com/Eli-Meyer/TagSeq_utilities) was used to process the RNASeq data. Briefly, the quality of raw sequences was assessed using ‘FastQC’v.0.11.9. Raw reads were filtered based on Phred per-base quality score (Q) ≥ 20 and a cumulative per-read low quality score (LQ) ≤ 10 using the QualFilterFastq.pl script with the parameters -m 20 and -x 10. Reads that passed this step were then depleted of homo-polymer repeats longer than 30 bp using HRFilterFastq.pl (parameters -n = 30), adapter sequences using BBduk.sh, part of BBMap v.35.85, and PCR duplicates using RemovePCRDups.pl. Non-template sequences introduced at the 5’ end of cDNA tags during Tag-Seq libraries preparation were trimmed using TagTrimmer.pl (parameters: -b 1 -e 8).

Quality-processed reads were re-evaluated for quality using FastQC (https://www.bioinformatics.babraham.ac.uk/projects/fastqc/) and then mapped using HISAT (Kim *et al*., 2015) to the *Sorghum bicolor* reference. SAMtools v.1.10 (Etherington *et al*., 2014) was used to convert the generated sequence alignment map (SAM) files into binary alignment mapping (BAM), and to sort and index the BAM files. Transcripts were quantified using StringTie v.2.1.1 (Pertea *et al*., 2014) to generate normalized counts in form of FPKM and TPM in addition to RAW counts for differential gene expression analysis.

### Identification of differentially expressed genes

Raw counts were transformed into counts per million (CPM) using the *cpm()* function in edgeR package (Robinson *et al*., 2010). Genes with a CPM greater than 1 were retained for further analysis. Principle coordinate analysis (PCoA) plots and heatmaps were generated using ggplot2 (Wickham, 2011) to determine the relatedness of the biological replicates.

Differential gene expression analysis was performed using DESeq2 Bioconductor R package following normalization using DESeq2 (Love *et al*., 2014) with a Wald test. A false discovery rate (FDR) cut-off of 0.05 was applied, and a log2 fold change cutoff of ≥2 to indicate upregulation and ≤-2 to indicate down-regulation. Differentially expressed genes (DEGs) were considered at each time point for each host in relation to the controls. Pathway enrichment analysis of DEGs was performed using the classical method and Fisher’s exact test with a P-value threshold of ≤0.05 using ShinyGO version 7.48 (Ge *et al*., 2019).

### Weighted gene co-expression network analysis

Weighted gene co-expression networks (WGCNA) were performed using WGCNA (v1.71) package in R (Langfelder and Horvath, 2008). The DESeq2 *dds* normalized counts were transposed and then passed on to WGCNA to construct co-expression modules using the automatic network construction function blockwiseModules with default settings. The power was set at 10 after determining the optimum threshold using the ‘*pickSoftThreshold*’ function. TOMType was signed. The correlation coefficient for module eigengenes was calculated using Pearson’s correlation and hubs were selected using *‘chooseTopHubInEachModule’* function. For visualization, modules were then exported to Cytoscape_3.9.1 (Shannon *et al*., 2003).

## Results

### Diverse phenotypes of *Striga* resistance in sorghum

We selected sorghum genotypes previously described in the study by Kavuluko *et al.* (2021). In the paper, we described N13 and IS9830 as *Striga* resistant based on inability of the parasite to penetrate the host and establish vascular connections. This resistance was attributed to fortification of the host cell wall by various polysaccharides. In the same study, IS14963 was characterized as resistant due to elicitation of an intense hypertensive reaction leading to programmed cell death at the interphase of the host and the parasite. Along with those genotypes, we also evaluate IS2814 and IS41724 whose mechanism of resistance had not been characterized prior. Our analysis showed that IS2814 had a resistance phenotype that blocked parasite ingression into the host endodermis. The parasite’s infectious organ – haustorium – did not contact the host vasculature in the resistant genotype IS2814. Instead, the parasite haustorium bypassed the host endodermis, and emerged from host root tissue at the distal end (Fig.1a). In IS41724 our analysis showed that programmed cell death at the interphase of the host and the parasite. A close-up of the parasite attaching the host shows a poorly developed parasite seedling tissue necrosis at the interphase of the parasite and the host. A transverse section through the haustorium affirmed the hypersensitive reaction as the main resistance mechanism in this genotype (Fig. 1b).

**Fig. 1.**
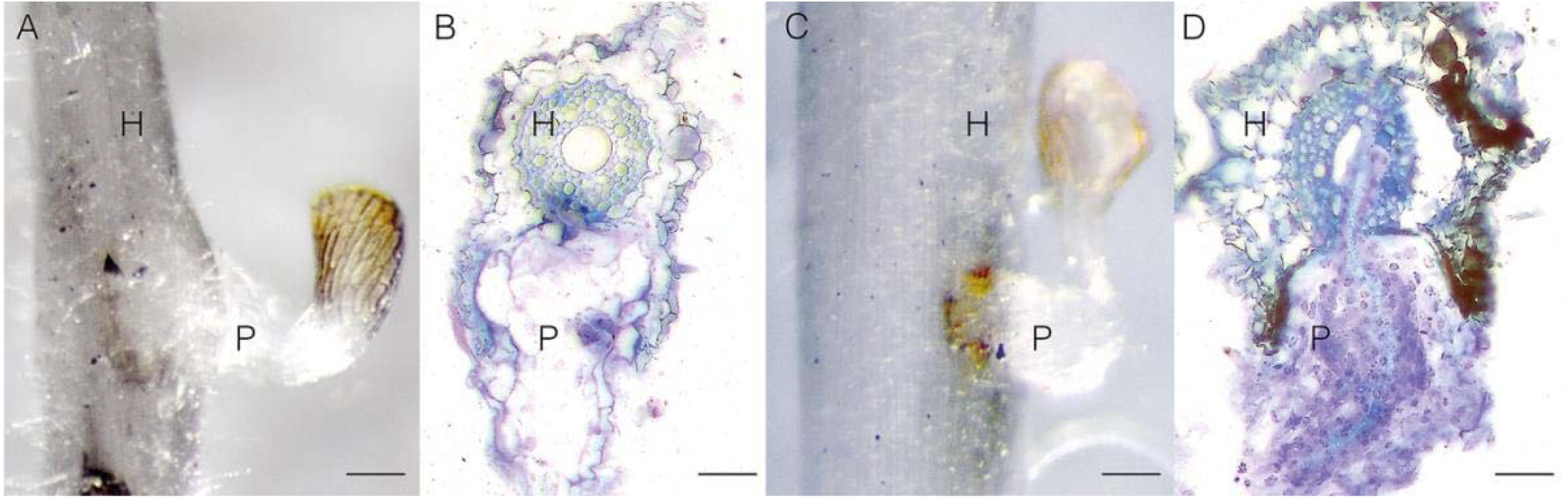
Contrasting phenotypes of *Striga* resistance: (A and B) Cell wall enhancement-based resistance in IS2814 showing parasite ingression blockage by cell wall barriers. (A) Close-up showing a poorly developed parasite. Scale bar = 5 mm. (B) A transverse section shows parasite tissue cycling around the host endodermis. (C and D) Hypersensitive reaction-based resistance in IS41724. (C) Parasite tissue starts a hypersensitive reaction at the interphase of the host and parasite. (C) Closeup shows a poorly developed parasite tissue and the beginning of necrosis. (D) Transverse sections show necrosis in areas of attempted penetration by the parasite. Scale bar for close-up images is 5 mm, and 0.5 mm for transverse section images. H = Host, P = Parasite.

These results show that *Striga* resistance in IS2814 and IS41724 is due to cell wall enhancement and elicitation of a hypersensitive reaction, respectively. In the end, the resistance phenotypes of sorghum used in the study were described as displaying either mechanical barrier resistance (N13, IS9830 and IS2814) or hypersensitive reaction (IS14963 and IS41724).

### Global transcriptome reprograming in sorghum upon *Striga* infection

To understand the global transcriptome reprograming that occurs following infection by *Striga,*and the underlying genetic causes of resistance for each genotype, we analyzed differentially expressed genes (DEGs) in each sorghum genotype at early 3 days after infection (dai) and late stages (9 dai). We found unique genes that were differentially expressed for each genotype suggesting a genotype dependent transcriptional activation of defense genes. We also found DEGs that were shared among various genotypes and even core genes that were shared by all genotypes at each stage, suggesting that some transcriptional responses were common and conserved in sorghum (Fig. 2).

**Fig. 2.**
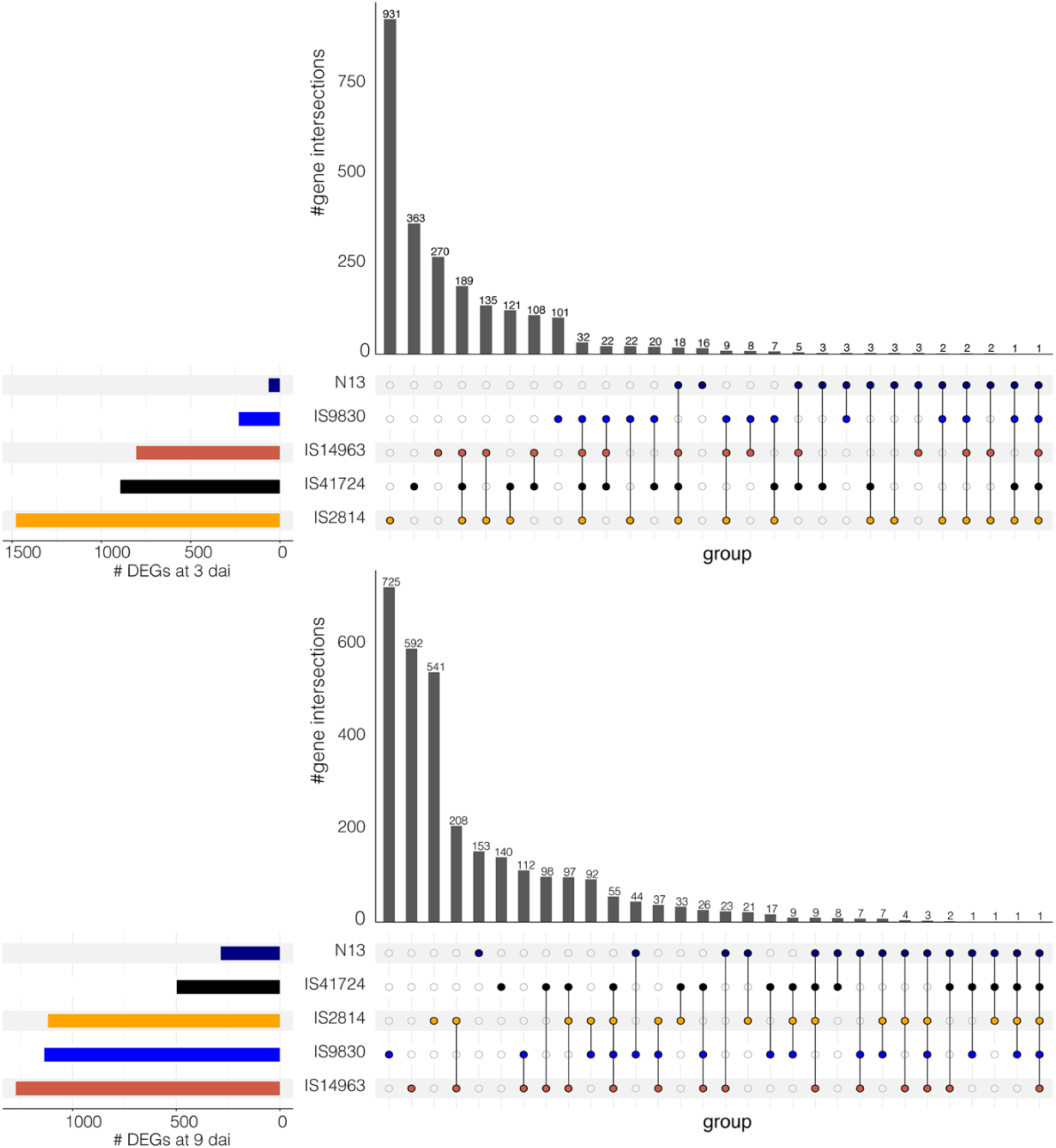
Transcriptional changes in sorghum upon *Striga* infection. Upset plots showing how different genotypes of sorghum respond to infection by *S. hermonthica* at 3 dai (upper panel) and 9 dai (lower panel). The plot compares the overlap of differentially expressed genes (DEGs) between comparisons with the colored horizontal bar graphs indicating the number of DEGs for each comparison, respectively. The black vertical bar graphs show the intersection size of DEGs and the colored dots represent contributing comparisons from the DE analysis.

### WGCNA clarifies phenotypes of *Striga* resistance in sorghum

To further resolve the complexity of resistance responses in the various sorghum phenotypes, we performed a weighted gene co-expression network analysis (WGCNA). This analysis clusters DEGs based on their expression levels in different samples and groups them as modules. Thus, WGCNA allowed us not only to identify modules associated with various modes of resistance in individual genotypes, but to also group sorghum genotypes showing similar gene expression patterns and identify critical genes for each regulatory network (Fig. 3).

**Fig. 3.**
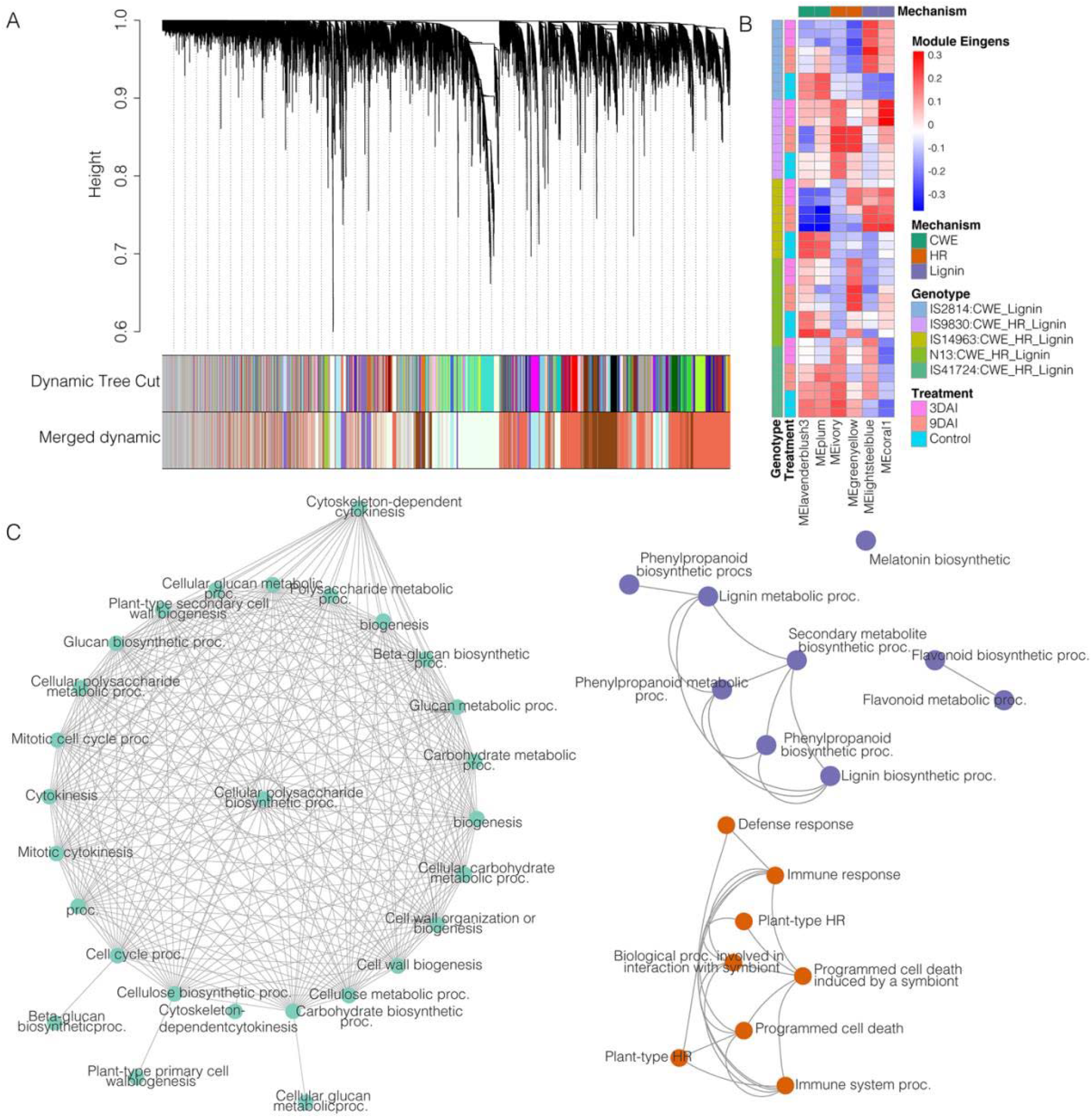
Weighted Gene Expression Network Analysis (WGCNA) of 5 *Striga* resistant sorghum transcriptomes. (A) A Dynamic cut tree of 43 module and 13 merged modules. (B) A heatmap showing clarification of mechanisms of resistance using WGCNA module Eigengene values. Six modules corresponded with three mechanisms of *Striga* resistance. Cell wall enhancement (CWE), Hypersensitive reaction (HR) and secondary metabolites including lignin biosynthesis. The heatmap also shows that WGCNA also partitioned genotypes according to modes or resistance. (C) Pathway interaction networks representing *Striga* resistance mechanisms: MEIavendablush3 representing with significant enrichment for primary and secondary cell wall reinforcements, most notably callose apposition though beta glucan biosynthesis. MEgreen-yellow module showing network processes for programmed cell death, and MEcorall showing enrichment for secondary metabolites biosynthesis including lignin and melatonin. Network nodes are color-coded according to the modes of resistance.

We found a total of 77 statistically significant modules (Dataset S1). A gene ontology (GO) enrichment analysis of the modules revealed that 6 modules were enriched for processes of: i) cell wall enhancement, ii) HR response and iii) secondary metabolites production including lignin biosynthesis (Fig. 3 and Fig. S2). Cell wall enhancement pathways were enriched in the MElavenderblush3 and MEplum modules for processes of, cell wall growth like cell wall biogenesis, cell wall organization and microtubule development. Additionally, the modules were enriched for the synthesis of cell wall polysaccharides, for example, beta-glucan and cellulose biosynthesis. Hypersensitive resistance mechanism pathways were in the MEivory and MEgreen-yellow. Here, processes of programmed cell death, hypersensitive response, and immune activation consistent with host-pathogen interactions were enriched in both modules, but more notably in MEivory. Finally, MElightsteel blue and MEcoral1 were enriched for pathways consistent with the production of secondary metabolites that enhance resistance against invading pathogens. In these modules, we found enrichment for upstream metabolites of lignin biosynthesis – the phenylpropanoid pathway as well as flavonoids. Notably, MEcoral1 was also enriched for melatonin biosynthesis. This finding was unexpected, but significant because, as it happens, melatonin increases lignification (Zhao *et al*., 2022).

We grouped genotypes showing similar molecular signatures of *Striga* resistance and found consistent clustering of genotypes and treatments according to module Eigengene values. This allowed us to assign resistance mechanisms for the various genotypes as follows: (i) IS2814 was strongly positively correlated with modules for cell wall enhancement and lignification, (ii) IS9830, IS14963, N13, and IS41724 were all correlated with the three modes of resistance (CWE, HR and lignification) albeit at different degrees (Fig. 3).

The final aspect of our WGCNA analysis was to identify transcriptional regulatory pathways and associated top network genes in the various individual and merged modules involved in *Striga* resistance. A list of annotated top genes in each module is provided in Table S1. We would like to highlight 3.

Firstly, Sobic.001G529600, in the green module. This gene encodes Glucan synthase-like 10 encodes (GSL10), a member of the Glucan Synthase-Like (GSL). GSLs are known to catalyze the synthesis of the cell wall component callose, which provides structural cell wall reinforcement against pathogen infection (Bacete *et al*., 2018). A closer look at the enriched pathways in the green module revealed significant enrichment for pathways involving innate immune response activating receptor signaling, immune response receptor transduction, and activation of immune responses – suggestive of a PAMP-induced callose deposition (Supplementary Figure 4). Consistent with callose deposition, the green-yellow network was enriched for beta glucan, xylan, and other carbohydrates that converged at the polysaccharides.

Secondly, the Sobic.004G321100 in the green-yellow module. This gene encodes a phosphoinositide phosphatase and forms an important component in the phosphoinositide signaling mediated defense network for both basal and systemic responses (Hung *et al*., 2014). Consistent with this view, we found significant enrichment of glycolipid and phospholipids biosynthetic and metabolic processes. Reinforcing our hypothesis, we found significant enrichment of cell death processes in the module (Fig. S2).

Thirdly, Sobic.006G280000, in the turquoise module which encodes the pathogenesis-related thaumatin like family of genes is consistent with activation of the basal immune response to pathogens. It is known that plants produce pathogenesis-related (PR) proteins that play key roles in plant disease-resistance responses, specialized in SAR (Hamamouch *et al*.,2011). Expectedly, pathways for responses to wounding and basal-triggered immunity were enriched in this module (Fig. S2).

### *Striga* activates multiple defense pathways in sorghum

To determine specific defense responses that *Striga* triggers in sorghum, we studied gene expression patterns of sorghum in the key pathogen detection, signaling and defense pathways that included, i) Plant pathogen interaction, ii) phenylpropanoid and iii) hormone signaling pathways based on Kyoto Encyclopedia of Genes and Genomes (KEGG) maps.

#### (i) *Striga* elicits both pathogen and effector triggered immunity in sorghum

In this pathway both PTI and ETI are activated leading to downstream defense processes (Fig. 4). On PTI, our results showed that in all genotypes CNGCs, involved in Ca^2+^ sensing were induced at all stages. But CDPK and RBo genes leading to HR responses were only induced in IS9830 and N13. CaM/CML and FLS2 that lead to cell wall enhancement and other disease response reactions including WRKY and PR1-dependent disease responses were induced only in IS9830. On ETI, resistance genes RIN4, RPMI and RPS2 and HSP2 were induced in all genotypes – but more notably in N13 and IS9830. RPS2 was also notably induced in IS41724 at 3dai. In all cases, ETI genes were not induced in IS2814. These expression patterns suggest that ETI responses are induced in N13, IS9830, IS14963 and IS41724 and that PTI is induced in all genotypes for upstream genes but more notably in N13 and IS9830 for downstream genes. This observation is consistent with module enrichment results that grouped N13 and IS9830 as displaying both cell wall-based resistance as well as hypersensitive responses. These results are also consistent with phenotypic observations that showed HR responses in IS41724.

**Fig. 4.**
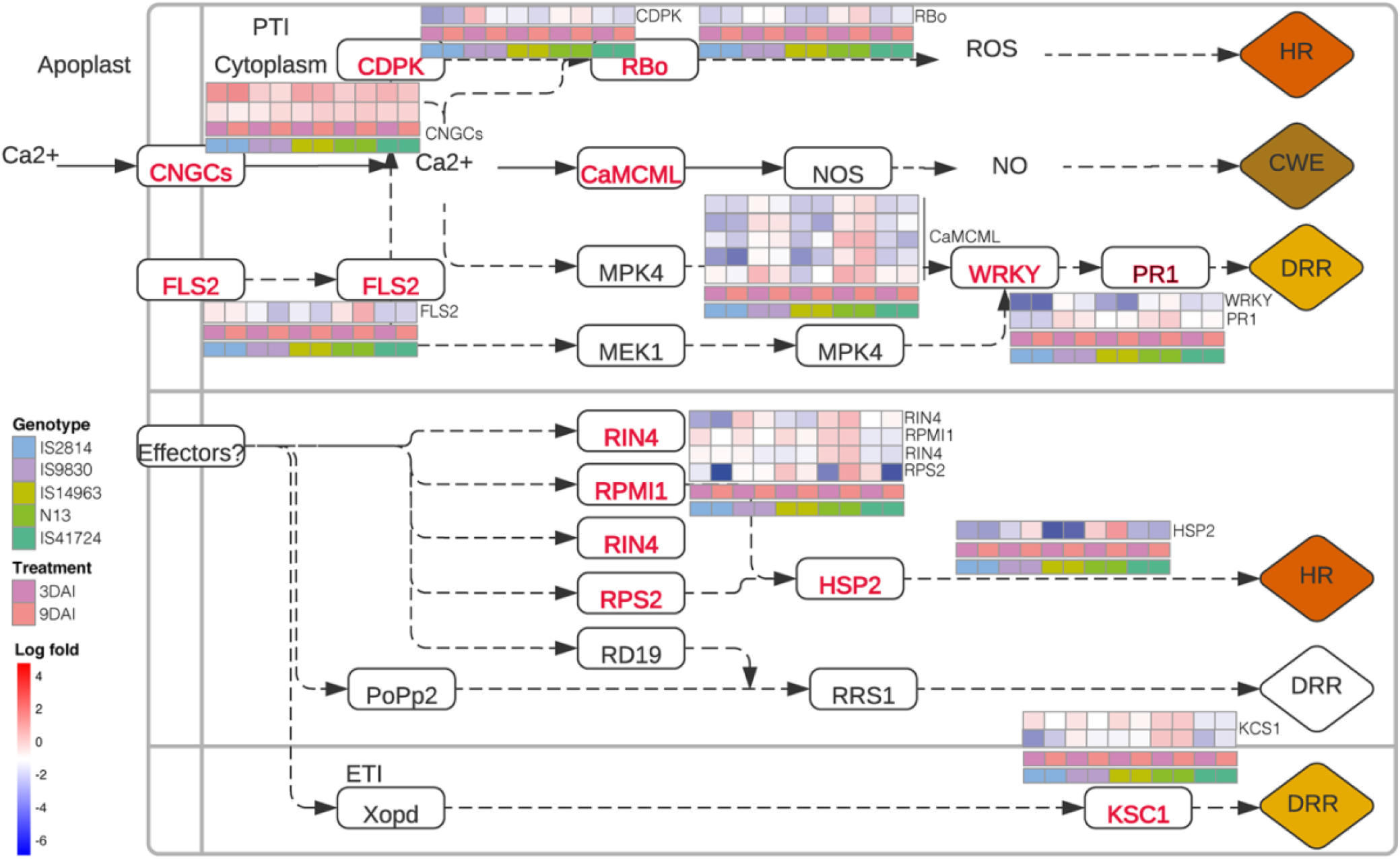
*Striga* elicits both pathogen and effector triggered immunity in sorghum. Gene expression patterns of sorghum genotypes in the plant-pathogen interaction Kyoto Encyclopedia of Genes and Genomes (KEGG) map. *Striga* induced pathogen and effector triggered immunity and showed differential gene expression patterns in the sorghum genotypes for various pathogen immune responses at varied degrees. In IS983Oand N13, genes were induced for hypersensitive responses (HR) responses as well as systemic acquired resistance (SAR) though the WRKY signaling pathway. The red-colored font of enzymes indicates the differentially expressed genes in response to *S. hermonthica* infestation. Red box denotes up-regulated expression while a blue box denotes a down-regulated expression.

#### (ii) Lignin biosynthesis pathway is critical against *Striga* resistance in sorghum

Next, we looked at the phenylpropanoid pathway that leads to lignin biosynthesis (Fig. 5a). Modules enriched for lignin biosynthesis (lightbluesteel and coral1) had the following annotated genes: one phenylalanine ammonia lyase (PAL), Ferulate 5-hydroxylase (F5H) Caffeic acid O-methyl transferase (COMT), Cinnamyl alcohol dehydrogenase (CAD) and 5 peroxidase genes. All these genes were significantly induced in IS2814, N13 and IS9830 in early and late stages of infection. The induction was more notable in IS2814 compared to N13 and IS9830. In comparison, induction levels of the genes remained lower in IS41724 and IS14963. These results reinforced WGCNA results that showed significant enrichment of lignin biosynthesis in IS2814, N13 and IS9830. To further examine lignin as a defense mechanism, we performed lignin staining assays on N13, IS41724 and IS14963. Consistent with gene expression results, lignin staining was highest in N13, moderate in IS41724 and almost undetectable in IS14963 (Fig 5b).

**Fig. 5.**
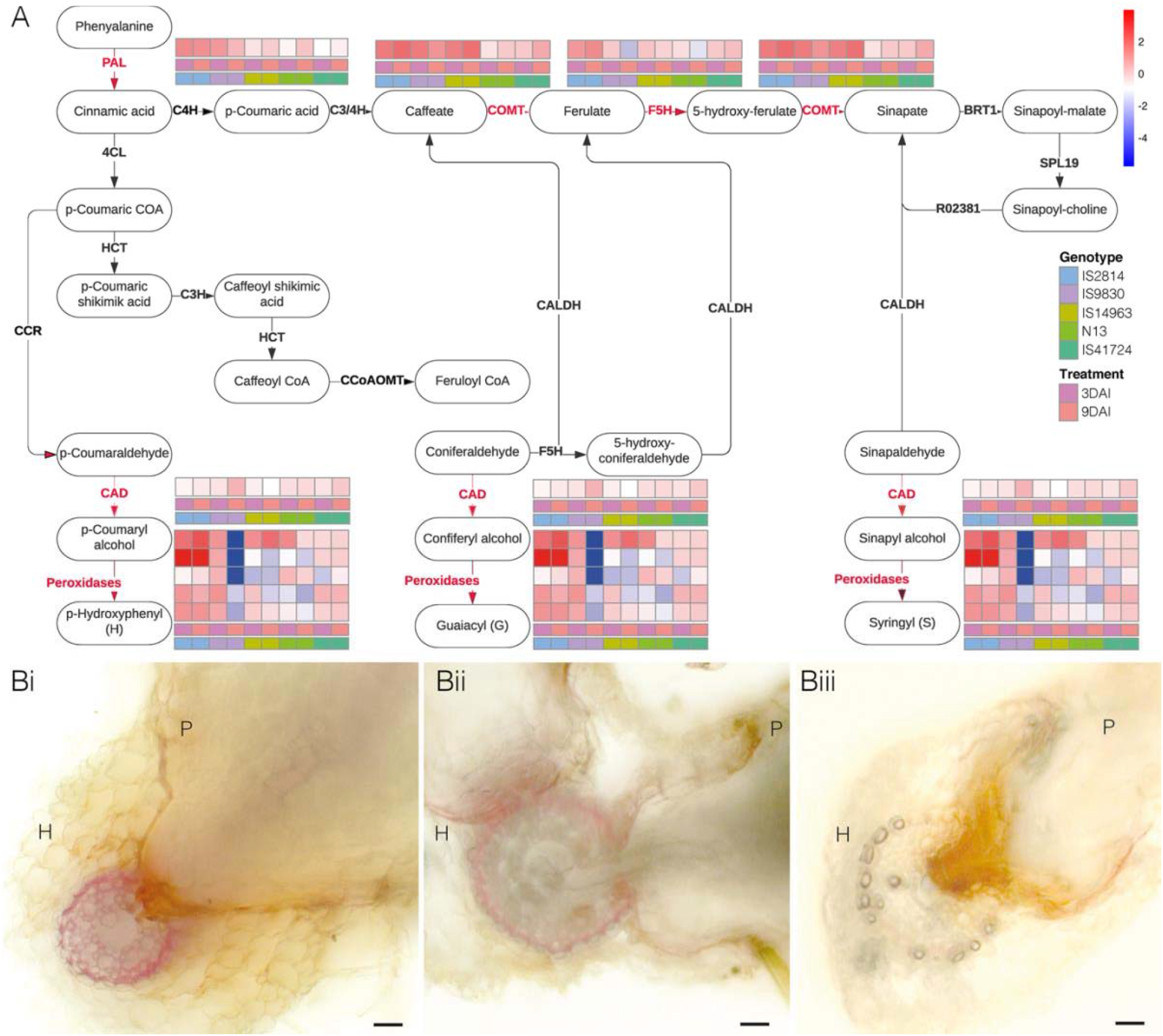
Lignification as a defense mechanism against *Striga* in sorghum. (A) Differential gene expression in the phenylpropanoid biosynthetic pathway leading to lignin biosynthesis. Key lignin genes were positively regulated in IS2814, and N13 compared to other genotypes. The red-colored font of enzymes indicates the differentially expressed genes in response to *S. hermonthica* infestation. Red box denotes up-regulated expression while a blue box denotes a down-regulated expression. (B) Histochemical staining using Phloroglucinol-HCI for lignin detection. Varying red color intensity in vibratome (3Oum) thick sections in: (Bi) N13, deepest coloration, (Bii) IS41724 moderate and faint in (Biii) IS14963. Scale bar is 0.1 mm; H = Host; P = Parasite.

#### (iii) *Striga* activates JA and SA immune responses in sorghum

Lastly, we examined the hormone signaling pathway focusing on jasmonic acid (JA) and SA signaling (Fig. 5). JA triggers the degradation of JASMONATE ZIM DOMAIN (JAZ) transcriptional repressor proteins leading to release of downstream transcription factors that in turn activate defense responses including programmed cell death. Correspondingly JA mediated defense, is followed by suppression of JAZ genes. Consistent with JA activation, JAZ expression levels were 4 to 8-fold lower in 9 JAZ proteins for the genotypes: IS2814, IS14963 and IS41724. In the other two genotypes, IS9830 and N13, JAZ levels remained the same. The levels of MYC2 were also unchanged. Interestingly, levels of SA associated hormones remained high in all genotypes. These results indicate that JA mediates genotype specific *Striga* resistance responses in genotypes IS2814, IS14963 and IS41724 exhibit *Striga* resistance in a JA dependent manner; and that *Striga* triggers a SAR in all the genotypes.

**Fig. 6.**
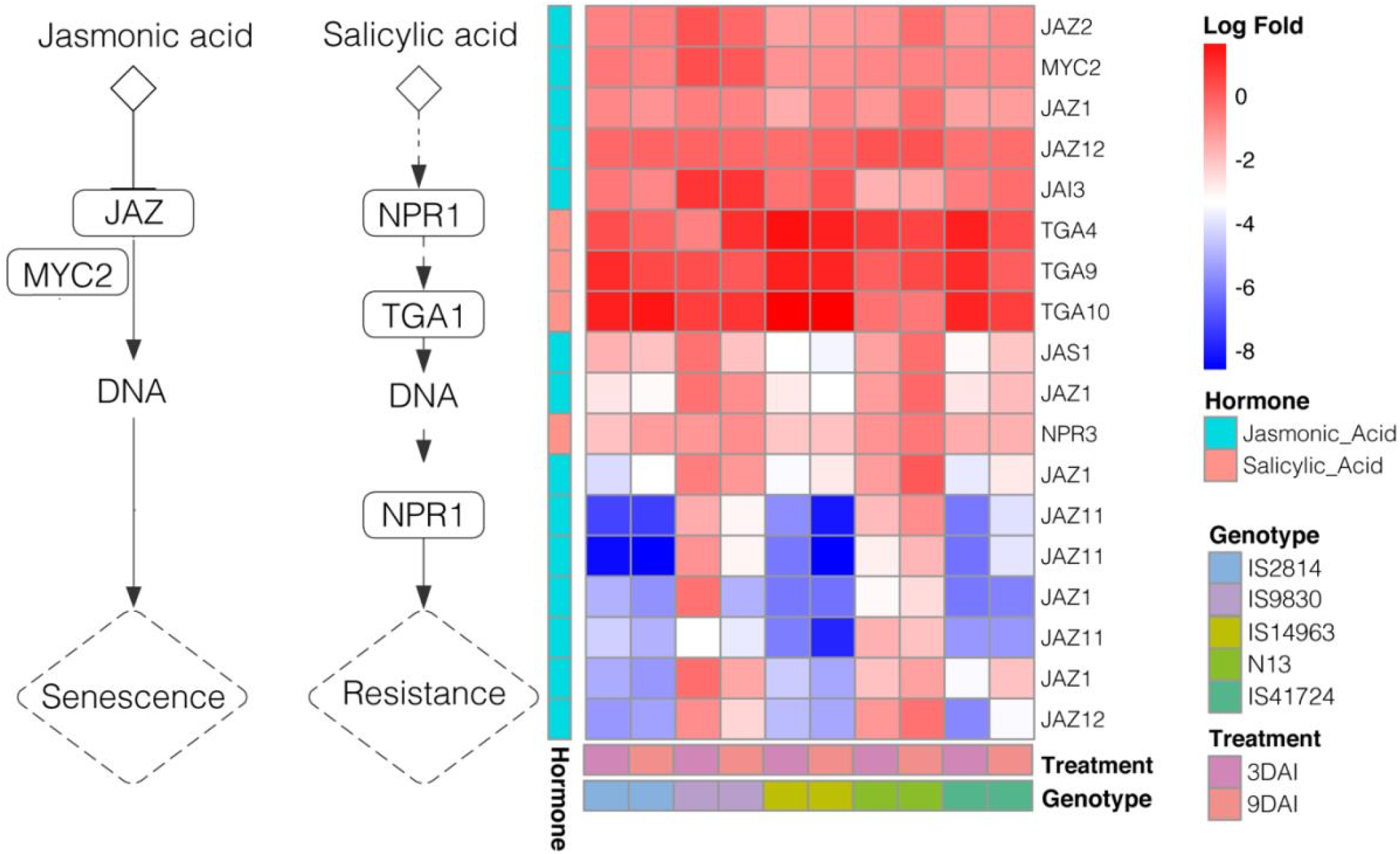
*Striga* activates both jasmonic acid (JA) salicylic acid (SA) downstream immune responses in sorghum. Differential activation ofJA and SA genes in different *Striga* genotypes. IS9830 and N13 showed a marked induction of JA responses. In contrast, SA responses were induced in all genotypes.

## Discussion

We sought to clarify genetic causes of different modes of *Striga* resistance in sorghum. We used 5 sorghum genotypes with distinct documented mechanisms or resistance. Three of the genotypes had mechanical resistance and two had HR responses. We found that sorghum displayed many differentially expressed genes for each of the genotypes, underscoring the fact that sorghum has a large genetic pool that can be exploited for improvement. Bearing in mind that each genotype had a different biological status: IS41724 and N13 are Dura, IS9830 and IS14963 Caudatum and IS2814 Durra-Caudatum sorghums (Kavuluko *et al*., 2021). And that, the genotypes were domesticated in different regions of Africa (Kavuluko *et al*., 2021). Unique sets of genes responding to infection by the parasite suggest independent adaptation and evolution of the resistance and shared genes show conservation of key pathways for resistance against the parasite.

### Do sorghum genotypes harbor distinct resistance mechanisms against *Striga*?

We used WGCNA to further decipher these complexities. In this analysis, reads are weighted based on similarity of gene expression and assigned modules based on this similarity. We found that WGCNA efficiently delineated mechanisms of *Striga* resistance and placed them in modules bearing common molecular signatures of resistance. From these groupings, we were able to classify the genotypes according to resistance mechanisms based on activated pathways. WGCNA grouping bore truth to earlier observations and provided new insights in sorghum resistance against *Striga*. For example, N13 has been described as having quantitative resistance based on cell wall enhancement traits (Maiti *et al*., 1984; Haussmann *et al*., 2004), in this study, IS2814 was shown to harbor mechanical barrier resistance and, IS41724 and IS14963 were shown to display HR resistance.

### Can WGCNA help pinpoint key *Striga* resistance genes in sorghum?

An additional feature of our WGCNA analysis was identification of top genes regulating module networks. We focused on three key genes based on their documented role in plant-pathogen interactions.

The first gene of interest GLS10 is a key regulator of callose biosynthesis. Callose is a β-(1,3)- D-glucan is widely distributed in higher plants. In addition to its role in normal growth and development, callose plays an important role in plant defense (Voigt, 2014). In Arabidopsis, deposition of callose or papillae, at sites of fungal penetration serves as an early response of host plants to microbial attack provides protection against entry of the fungus (Jacobs *et al*., 2003). Because the green module network was also enriched with immune responses, we attributed this to a PAMP induced, GLS callose deposition as earlier described (Jacobs *et al*., 2003). This defense response is characterized by formation of calloses rich cell wall appositions called papillae (Jones and Dangl, 2006; Schwessinger and Ronald, 2012). Because of the earlier documented role of papillae in stopping penetration of fungal hyphae in barley (Chowdhury *et al*., 2014), it is not far-fetched to speculate that callose apposition may make the difference between successful and failed *Striga* penetration attempt. Indeed, overexpression of GSL5/PMR4 leads to enhanced papillary callose deposition and complete penetration resistance to powdery mildew, showing that cell wall carbohydrates can contribute to penetration resistance (Ellinger *et al*., 2013).

The second interesting gene, Sobic.004G321100 in the green-yellow module encodes a phosphoinositide phosphatase and forms an important component in the phosphoinositide signaling mediated defense network for both basal and systemic responses (Hung *et al*.,2014). Therefore, identification of this gene was indicative of a phosphoinositide signaling for basal and systemic response following *Striga* infection. The membrane-associated phospholipids along with the soluble inositol phosphates (collectively known as phosphoinositides) are present in all eukaryotic cells and are implicated in plant responses to many environmental stimuli [reviewed in (Stevenson *et al*., 2000)]. The phosphoinositide pathway and inositol-1,4,5-triphosphate (InsP3) have been implicated in mediating both basal defense and systemic acquired resistance responses (Hung *et al*., 2014). The study showed that Flagellin induced Ca^2+^ release as well as the expressions of some flg22 responsive genes were attenuated in the InsP 5-ptase plants, which had reduced InsP3, making them more susceptible. Additionally, InsP 5-ptase plants were more susceptible to virulent and avirulent strains of *Pseudomonas syringae pv. tomato (Pst)* DC3000 and had lower basal salicylic acid (SA) levels and the induction of SAR in systemic leaves was reduced and delayed. Plausibly, phosphoinositide signaling is one component of the plant defense network and is involved in both basal and systemic responses. Affirmation of this process can also be drawn from the significant enrichment of cell death processes in this module.

And the third, in the turquoise module the gene Sobic.006G280000, which encodes the pathogenesis related thaumatin like family of genes. Production of pathogenesis related proteins (PR) represents the hallmark of SAR is in plants (Hamamouch *et al*., 2011). Specifically, thaumamatin like proteins (TLPs) can rapidly accumulate to high levels in response to biotic or abiotic stress and exhibit antifungal activity in various plant species (Petre *et al*., 2011). Expectedly, over-expression of TLPs can induce resistance against pathogens. For example, over expression of TLP has been shown to increase resistance against fungal pathogens in tomatoes (Liu *et al*., 2012) and potato (Acharya *et al*., 2013). Based on the gene expression patterns observed in our studies, we infer that TLP would be good candidates to impart *Striga* resistance. Support for this hypothesis can be drawn from gene expression studies that have shown up regulation of PR genes in the resistant rice variety (Nipponbare) relative to the susceptible IAC (Swarbrick *et al*., 2008). In the Swarbrick *et al*. (2008) report, genes encoding endochitinases (PR-3), glucanases (PR-2) and thaumatin-like proteins (PR-5) were up regulated in the resistant rice variety. Furthermore, a subsequent similar study in cowpea-*Striga* interactions showed that levels of PR5 were dramatically upregulated in the resistant variety (Huang *et al*., 2012).

### *Striga* elicits both pathogen and effector triggered immunity in sorghum

Individual gene expression further supported activation of PTI and ETI for various modes of resistance. At the PTI level, there was upregulation of HR response genes – CNGS, CDPK and RBO and SAR associated genes, WRKY and PR1 in IS9830 and N13 consistent with the modes of resistance of these genotypes. At the ETI level, resistance genes RIN4, RPMI, RPS2 as well as HSP and KSC were up regulated in IS9830 and N13. Downstream pathways of ETI and PTI led to activation of defense processes such as cell wall enhancement that were also consistent with the modes of resistance for the genotypes. The concept of an active PTI triggered by *Striga* infection, raised the question as to what sorghum recognizes as PAMP. We hypothesized that products of cell wall degradation created upon *Striga* infection act as elicitors of PTI akin to damage associated pathogen patterns (DAMPS) previously described (Souza *et al*., 2017) but also plausibly, *Striga* has PAMPS. The later hypothesis is reinforced by identification of a cell wall-localized glycine-rich protein that acts as a PAMP in the parasitic plant *Cuscuta reflexa* (Slaby *et al*., 2021).

### Lignin biosynthesis pathway is critical against *Striga* resistance in sorghum

We examined the phenylpropanoid pathway that leads to lignin biosynthesis and found up-regulation of key lignin biosynthesis genes – PAL, COMPT, FSH, CAD and peroxidases – in IS2814, IS9830 and N13 that exhibited cell wall-based resistance modes. In concert, an increased expression of lignin genes has been reported in cowpea infected with *S. gesnerioides* (Huang *et al*., 2012) and *S. hermonthica* rice interactions (Swarbrick *et al*., 2008; Mutuku *et al*., 2019) suggesting that lignification is a critical mechanism of defense employed by hosts to stop *Striga* parasitism.

In the defense signaling pathway, the most differentially induced genes were those involved in JA signaling and response, particularly JAZ – and this expression patterns corresponded with the resistance modes. Notably, in IS9830 and N13, JAZ genes were notably induced. In striking contrast, SA response genes, NP3 and TGA were induced in all genotypes suggesting active JA and SA signaling in the sorghum genotypes. In a previous study (Hiraoka and Sugimoto, 2008) of differential transcriptional response of susceptible and moderately resistant sorghum to *S. hermonthica*, the authors found that *Striga* parasitism induced JA-responsive genes and suppressed SA-responsive genes in the roots of a highly susceptible cultivar and suggested that susceptible hosts recognize *Striga* parasitism as wounding stress rather than microbial stress. In contrast, they found that in the roots of a moderately resistant sorghum cultivar, both SA- and JA-responsive genes were induced suggesting that resistance involved pathways associated with both wounding and pathogen challenge. In *Striga*-rice interaction, Mutuku *et al*. (2015) reported that both JA and SA pathways were induced, but the induction of the JA pathway preceded that of the SA pathway and that foliar application of JA resulted in higher resistance. These seemingly contrasting findings suggest that JA and SA signaling pathways are host-pathogen specific, but such a hypothesis will require further investigation.

## Conclusions

Taken together, our analysis of sorghum resistance to *S. hermonthica* is suggestive of diverse mechanisms for surveillance, signaling and defense. We identified important candidate genes that impinge on these processes that eventually impart *Striga* resistance. This makes future sorghum improvement for *Striga* resistance feasible and possible through breeding or genetic modification approaches.

## Abbreviations

CWE: Cell wall enhancement
DEGs: differentially expressed genes
ETI: effector triggered immunity
HR: hypersensitive response
PTI: pathogen triggered immunity
SAR: systemic acquired resistance
SSA: sub-Saharan Africa
TLPs: thaumamatin like proteins
WGCNA: weighted gene co-expression network analysis

## Supplementary data

The following supplementary data are available at JXB online.

Table S1: Top-network genes for single and merged modules

Fig. S1. Network topology for soft threshold power

Fig. S2. Enrichment analysis for key network genes

Fig. S3. Sequence alignment of CDKE homologues. Dataset S1. A spreadsheet of module Eigengene values

Dataset S1. A spreadsheet of module Eigengene values

Dataset S2. A spreadsheet of module classifications, gene ids and annotations

## Acknowledgement

We would like to acknowledge the International Crops Research Institute for the Semi-Arid Tropics (ICRISAT) through Dr. Eric Manyasa for availing the sorghum germplasm used in this project.

## Author Contributions

Conceptualization: SR, EB, SM. Methodology: SM. Sequencing: EB, BW, A.W. Data curation and analysis: SM, FM, EB, SR. Validation: OD, SR, SW. Writing original draft: SM, SR. Manuscript Editing: EB, EA, SW. All authors reviewed and approved the final Manuscript.

## Data availability

Raw reads from transcriptome sequencing have been deposited in the National Center for Biotechnology Information (NCBI) Sequence Read Archive under BioProject accession number PRJNA906112.

